# Evaluating cell type annotations in single-cell omics in the absence of ground truth

**DOI:** 10.64898/2026.06.10.731285

**Authors:** Josep Garnica, Massimo Andreatta, Santiago J. Carmona

## Abstract

Accurate cell type annotation is essential for single-cell transcriptomics, directly shaping downstream analyses and biological interpretations. Yet, objective evaluation of annotation quality remains a major challenge. Here, we argue that a cell type or cell state label has practical utility only if it captures a molecular pattern that is reproducible across biological replicates. Based on this principle, we introduce inter-sample consistency (ISC), a quantitative framework to assess annotation quality in single-cell RNA-seq datasets. Unlike existing cluster validation approaches, ISC distinguishes annotations that generalize across samples and individuals from those driven by technical or unwanted variation, thereby providing principled criteria for annotation quality and transferability. When applied to published single-cell atlases, ISC reveals widespread reproducibility gaps and provides actionable guidance for repairing inconsistent annotations. Notably, ISC enables benchmarking of automated cell type annotation tools even when ground-truth labels are unavailable, providing interpretable metrics to guide their development and evaluation. Implemented as the scTypeEval Bioconductor package, this framework offers a broadly applicable resource for evaluating and improving cell type annotations in single-cell RNA-seq experiments.

## Introduction

A central goal of single-cell RNA sequencing (scRNA-seq) is to characterize the cellular composition of tissues by grouping cells with similar transcriptional profiles into discrete categories, commonly referred to as cell types or cell states. Accurate annotation of these categories is essential to the biological interpretation of scRNA-seq data^1–3^. The most widely used approach to cell type annotation relies on a manual, iterative workflow involving unsupervised cell clustering, inter-cluster differential gene expression analysis, and expert interpretation of marker genes^4–7^. This process is time-consuming, inherently subjective, and often inconsistent across studies, thereby limiting reproducibility and cross-dataset comparisons. Automated cell type classifiers offer faster and more standardized alternatives, but their performance is constrained by the availability and relevance of reference datasets for the tissue, condition, or disease context of interest^8,9^.

Regardless of the annotation strategy, objective assessment of annotation quality remains a central challenge. Benchmarking of cell type annotation has primarily relied on three classes of metrics: (i) agreement with reference annotations (e.g. using the adjusted rand index), (ii) marker-based consistency metrics, and (iii) reference-agnostic measures of cluster compactness, separability, and stability, e.g. silhouette coefficient, Local Inverse Simpson’s Index (LISI), k-nearest-neighbor (KNN) purity, or Ratio of Global Unshifted Entropy (ROGUE)^10–14^.

A fundamental limitation of approaches (i) and (ii) is their dependency on previously manually annotated datasets or curated marker sets, which can introduce systematic biases, particularly given the absence of gold standards for cell type definitions. While for coarse-grained cell type annotations (e.g. broad immune lineages^15^) there is some general agreement among authors, there is substantially less consensus for the fine-grained cell type definitions typically required for detailed biological interpretation^3,16–19^. Internal validation metrics in category (iii) are reference-agnostic and do not require prior cell type knowledge. However, these metrics assess internal transcriptional coherence and robustness within a single or aggregated dataset and do not explicitly account for inter-sample variability. Thus, they cannot distinguish clusters that are consistently recovered across samples from those driven by a subset of samples or dominated by sample-specific effects.

In practice, researchers often cite the recurrence of annotated cell identities across samples or individuals as evidence of biological relevance (e.g. “detected in X/Y patients”)^18,20–22^. While high recurrence can support cell type definitions and should be considered, limited recurrence across samples alone does not invalidate a cell type annotation. Rare or context-specific cell types may be biologically meaningful and analytically valuable, provided they exhibit well-defined and distinctive transcriptional features across the subset of samples in which they occur. In contrast, cell type definitions that lack coherent, shared features across samples offer limited utility, even if they are nominally present in many samples.

Despite its importance, this concept – hereafter referred to as “inter-sample consistency” (ISC) – has not been formalized for assessing cell type annotations. This work introduces a set of novel metrics that explicitly quantify ISC, systematically assessing their performance and robustness to factors such as batch effects and annotation granularity. We demonstrate the utility of the ISC framework as a diagnostic tool for evaluating cell type annotations and for benchmarking cell type classifiers in the absence of ground truth. To facilitate adoption, we provide scTypeEval, a Bioconductor package implementing ISC-based methods for assessment and refinement of cell type annotations in single-cell omics data.

## Results

### Cell type expression profiles are frequently inconsistent across samples

To illustrate the potential of ISC to assess cell type annotation quality, we evaluated three published scRNA-seq datasets with expert-curated annotations, spanning different tumor types and healthy tissues^23–25^ (Fig. 1 and Extended Data Fig. 1a). Each dataset comprised samples from at least 12 donors, generated using a consistent experimental protocol and preprocessing pipeline within the same laboratories, therefore minimizing technical variability and batch effects. We constructed pseudobulk gene expression profiles by aggregating cells of the same reported cell type within the same sample of origin, yielding one transcriptomic profile for each combination of cell type and sample. Principal components analysis (PCA) of these pseudobulk profiles revealed heterogeneous patterns across datasets. In the Joanito dataset^24^, profiles corresponding to the same cell type clustered tightly across donors and were well separated from other cell types, indicating high inter-sample consistency (Fig. 1a). This is expected for low-granularity cell type annotation, i.e. when only highly divergent cell lineages are distinguished. In contrast, the Sikkema^23^ and Chu^25^ datasets – annotated at finer granularity – exhibited substantial donor-to-donor variability for several annotated cell types, such that profiles from the same cell type often overlapped with those of other cell types, indicating lower inter-sample consistency (Fig. 1b and Extended Data Fig. 1a).

**Figure 1.**
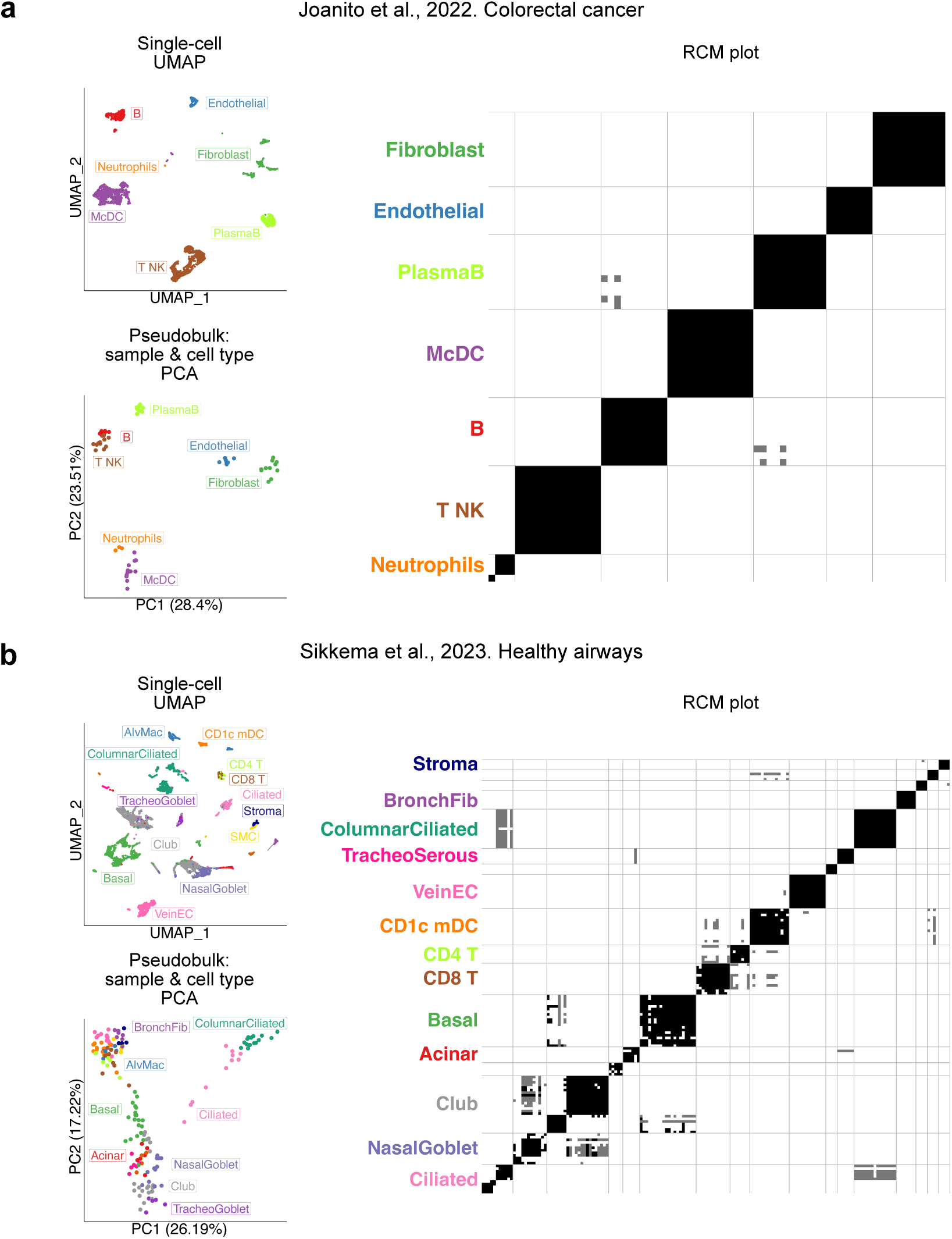
Inter-sample consistency of expert-curated cell type annotations in colorectal cancer and healthy airway scRNA-seq datasets. Two published datasets with expert-curated annotations are shown: Joanito et al. 2022^24^ (13 donors; colorectal cancer) (**a**) and Sikkema et al. 2023^23^ (22 donors; healthy airways; originally from Deprez et al. 2019^26^) (**b**). For each dataset, the left panels depict complementary views of the data: the upper panels show UMAP embeddings of single cells colored by annotated cell type, illustrating the structure of the data at single-cell resolution; the lower panels show PCA projections of pseudobulk gene expression profiles, generated by aggregating all cells of a given annotated cell type within each donor, yielding one profile per cell type and sample pair. The right panels display reciprocal-classification match (RCM) plots, summarizing pairwise comparisons of annotated cell types across all sample pairs within each dataset. For each pair of samples, classifiers were trained in both directions on pseudobulk profiles. Each tile represents the comparison of cell types between two samples and is encoded as follows: black, reciprocal best match in both directions; grey, one-directional match only, consistent with absence or non-recovery of the corresponding cell type in the reciprocal sample; white, no match. Symmetrically, each row and column represent a cell type from one sample.

To quantify these observations, we applied a “reciprocal classification match” (RCM) approach (see Methods). Briefly, for each pair of samples within a dataset, a cell type classifier was trained on cell type profiles from one sample and used to predict cell type labels in the other sample. The two samples were swapped and the procedure repeated. A cell type was considered consistent between a given sample pair only if it constituted a mutual best match under this reciprocal classification scheme; otherwise, it was considered inconsistent. Pairwise reciprocal best-hit relationships were visualized as “reciprocal classification match” plots. RCM plots revealed compact diagonal blocks in the Joanito dataset, indicating high inter-sample consistency across all annotated cell types, except for neutrophils (Fig. 1a). In contrast, the Sikkema dataset showed multiple off-diagonal-block matches and missing diagonal entries for several respiratory epithelial populations, including club cells, pointing to inconsistencies among closely related cell types (Fig. 1b). In the Chu dataset, regulatory T cells exhibited high inter-sample consistency, whereas CD8^+^ T central memory, CD4^+^ and CD8^+^ stressed T cells, and CD4^+^ naive T cells showed poor inter-sample consistency, in this case without strong similarity to other annotated cell types (Extended Data Fig. 1a).

To assess whether commonly used internal validation metrics could capture the ISC patterns revealed by RCM, we compared the resulting ISC scores to a reference-agnostic cluster quality measure, the LISI score^10^. This metric was computed at single-cell resolution and subsequently aggregated at the cell type level (cLISI), enabling direct comparison with ISC scores derived from pseudobulk reciprocal classification. ISC was quantified for each cell type as the proportion of sample pairs in which reciprocal best-hit relationships were satisfied. Across all datasets, we observed a weak and inconsistent concordance between ISC and cLISI (Extended Data Fig. 1b).

Together, these analyses demonstrate that expert-curated cell type annotations can exhibit markedly different levels of inter-sample consistency that are not captured by standard internal validation metrics, underscoring the need for systematic, sample-aware evaluation metrics.

### Defining cell type inter-sample consistency metrics and evaluation criteria

We devised a panel of ISC metrics based on commonly used cluster validation approaches, including average silhouette width-based measures (hereafter silhouette and 2-label silhouette), graph-based neighborhood purity, clustering agreement, and nearest-medoid agreement^27–29^. While these cluster validation metrics are commonly used for cell-level compactness and separability within individual datasets, they could be adapted to account for variability across datasets or samples (see Methods).

All ISC metrics take as input a cell-type by cell-type dissimilarity matrix, in which expression profiles are compared across samples for each cell type. These matrices can be computed in two ways: from single-cell distributions – e.g. using the Wasserstein distance – or from aggregated pseudobulk profiles, using measures such as Euclidean distance, cosine similarity, or reciprocal-classification-based dissimilarities. For reciprocal classification, a classifier is trained bidirectionally for each sample pair, and the resulting reciprocal predictions are encoded as binary-match-based or score-based dissimilarities (Fig. 2a, see Methods).

**Figure 2.**
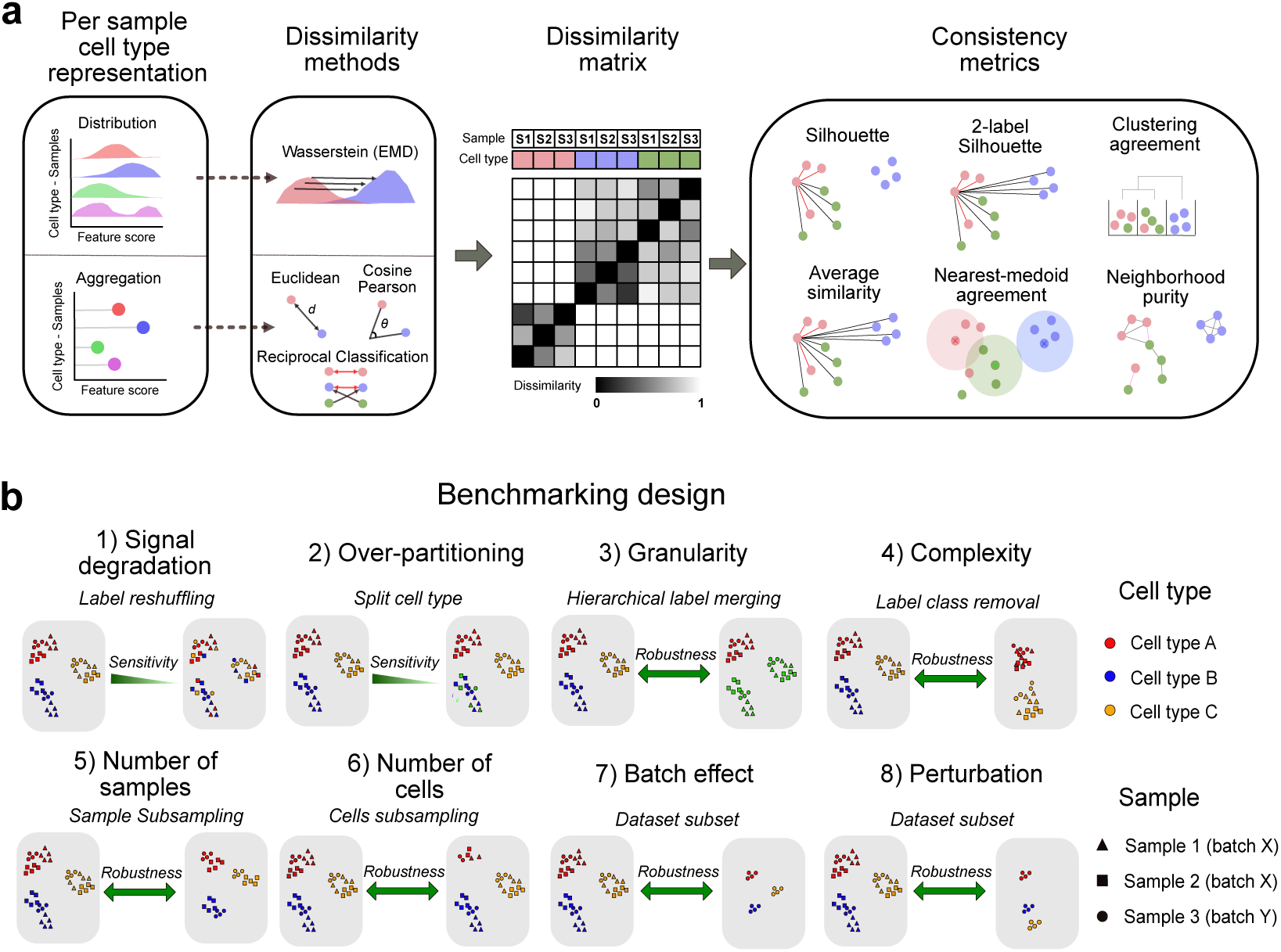
Definition and benchmarking of inter-sample consistency (ISC) metrics. (**a**) Overview of the ISC framework. Cell types are represented per sample either as single-cell expression distributions or as aggregated pseudobulk profiles. Pairwise dissimilarities between all cell type-sample combinations are computed using distribution-based distances (Wasserstein/EMD) or pseudobulk-based measures (Euclidean distance, cosine similarity, Pearson correlation, or reciprocal classification-based dissimilarities). These dissimilarities define a cell type-by-cell type matrix spanning all samples, which serves as input to sample-aware consistency metrics, including silhouette, 2-label silhouette, nearest-medoid agreement, clustering agreement, and neighborhood purity. Each metric yields a normalized consistency score per cell type, with higher values indicating greater inter-sample reproducibility. (**b**) Benchmarking design used to evaluate ISC metric behavior across eight criteria. Semi-synthetic perturbations were applied to 31 real scRNA-seq datasets to assess: (1) sensitivity to cell type signal degradation via progressive label reshuffling; (2) sensitivity to cell type over-partitioning by splitting a cell type into closely related subtypes; (3) robustness to annotation granularity through hierarchical label merging; (4) robustness to cellular complexity by removing subsets of cell types; (5) robustness to the number of samples via sample subsampling; (6) robustness to the number of cells per cell type via cell subsampling; (7) robustness to batch effects by comparing within-batch and pooled analyses; and (8) robustness to biological perturbations by comparing condition-specific and combined datasets.

Each ISC metric returns a scalar summary per cell type, with higher values indicating stronger inter-sample consistency. This formulation ensures that ISC metrics explicitly incorporate donor-to-donor variability and provide a quantitative framework to evaluate the reproducibility of cell type definitions across multiple samples. To systematically assess the behavior and utility of different ISC metrics, we defined eight evaluation criteria and corresponding tasks (Fig. 2b; see Methods):

(1) sensitivity to cell type signal degradation, evaluated by progressively disrupting cell type structure (compactness) in semi-synthetic datasets;
(2) sensitivity to cell type over-partitioning, assessed by artificially splitting cell populations into subpopulations;
(3) robustness to cell type annotation granularity, measuring consistency of ISC scores for coarse-versus fine-grained annotations;
(4) robustness to cellular complexity, comparing ISC behavior in tissues with low versus high inter-cell-type variability;
(5, 6) robustness to dataset size, quantified by varying the number of samples and the number of cells per cell type;
(7) robustness to batch effects, quantifying ISC stability in the presence of systematic technical differences;
(8) robustness to biological perturbations, evaluating ISC responses to systematic differences induced by biological conditions or experimental perturbations.

For each evaluation task, metric performance was quantified using a normalized “task performance score” ranging from 0 (worst performance) to 1 (best performance), enabling direct comparison across tasks (see Methods).

### Benchmarking inter-sample cell type consistency metrics

We benchmarked six ISC metrics combined with six dissimilarity functions across 31 scRNA-seq datasets from 17 studies spanning six biological systems: blood, colorectal cancer, melanoma, breast cancer, basal cell carcinoma, and respiratory tissue (Supplementary Table 1). Each dataset corresponded to a single batch and a relatively homogeneous biological condition, allowing us to evaluate metric behavior with limited confounding from strong batch or condition structure. For each of the eight evaluation tasks (Fig. 2b), these baseline datasets were perturbed or combined to generate semi-synthetic datasets with controlled changes in annotation quality, dataset composition, batch effects, and biological heterogeneity (see Methods).

A global ranking that integrates scores across tasks highlighted silhouette with RCM as the top-ranking ISC metric/dissimilarity function combination, followed by 2-label silhouette/RCM, and silhouette/pseudobulk cosine dissimilarity (Fig. 3a-b and Supplementary Table 2). This indicates that silhouette-based ISC metrics provide the best overall balance between sensitivity and robustness, particularly when paired with RCM. Pseudobulk-based methods were also markedly more efficient than distribution-based distances, typically running in under one second per dataset and using less than 100 MB of memory. Although Wasserstein distance performed well in some tasks, its substantially higher runtime and memory costs were not offset by improved overall benchmark performance (Fig. 3a).

**Figure 3.**
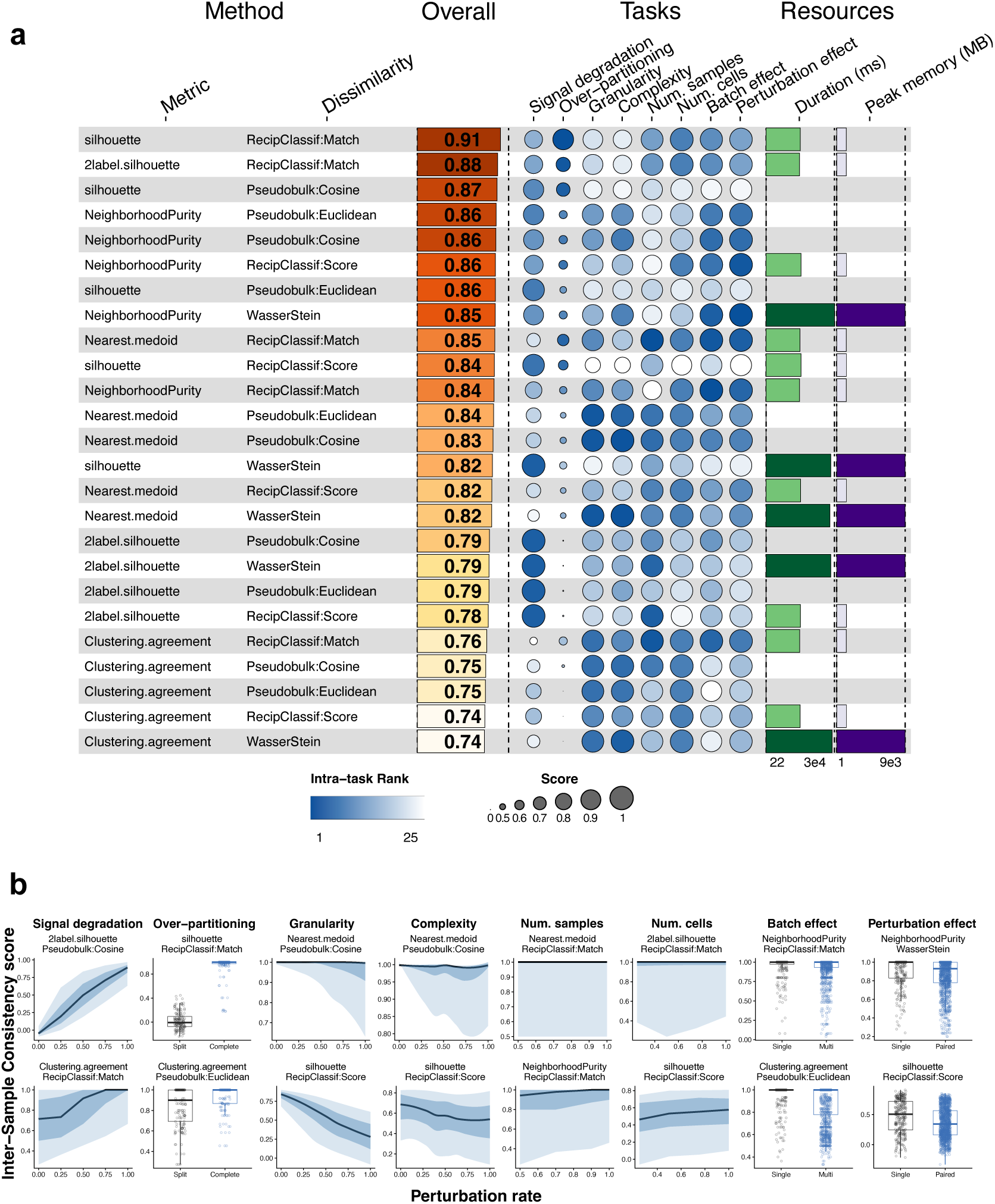
Comprehensive benchmarking of inter-sample consistency metrics across annotation quality and dataset complexity perturbations. (**a**) Heatmap summarizing the performance of 25 method combinations (5 inter-sample consistency metrics with 5 dissimilarity functions) across eight evaluation tasks. Average similarity metric and Pseudobulk:Pearson dissimilarity were removed from visualization, given their high correlation with 2-label silhouette and Pseudobulk:Cosine, respectively. Each row represents a method combination ordered by overall weighted score (left column). Task columns (middle) show median scores as colored circles, with circle size proportional to score magnitude (0-1 scale; legend) and color representing within-task rank (dark blue = best, white = worst). Resource columns (right) display median computational time (ms) and peak memory usage (MB) as colored bars, with darker shades indicating higher resource consumption. Overall scores represent weighted averages (Task 1: 20%, Task 2: 20%, Tasks 3-8: 10% each). n = 31 baseline datasets from 17 independent studies. (**b**) Method-specific responses to perturbation intensity across the eight evaluation tasks. Each column represents one task (labeled at top); rows show the best-performing (top) and worst-performing (bottom) method for that task. Y-axis: inter-sample consistency score. X-axis: perturbation rate or condition. Continuous perturbations (Tasks 1, 3-6) are displayed as line plots with ribbons showing 5th–95th percentile (light blue) and 25th–75th percentile (dark blue) ranges across all cell types and datasets. Two-level perturbations (Tasks 2, 7-8) are shown as box plots with jittered points comparing baseline (Split/Single) versus perturbed (Complete/Multi/Paired) conditions. Task 2 (Over-partitioning): Split = artificially split cell type, Complete = original annotation. Tasks 7-8: Single = within-condition consistency, Multi/Paired = cross-condition consistency.

To disentangle the contributions of the scoring metric and dissimilarity representation, we decomposed benchmark performance into marginal effects of each component (Extended Data Fig. 2a and Supplementary Table 2). 2-label silhouette contributed most strongly to task 1, indicating highest sensitivity to signal degradation, whereas silhouette contributed most strongly to task 2, indicating highest sensitivity to unsupported over-partitioning. At the dissimilarity level, RCM performed best for over-partitioning detection (task 2) and for robustness to batch and condition shifts (tasks 7 and 8), whereas pseudobulk cosine and Euclidean dissimilarities performed best for signal degradation detection (task 1). These results define two complementary dimensions of ISC: (i) task 1-oriented – best captured by 2-label silhouette with pseudobulk cosine dissimilarity, and (ii) task 2-oriented – best captured by silhouette with RCM dissimilarity. Most methods were robust to annotation granularity, cellular complexity, sample number and cells per cell type, with neighborhood purity as the main exception when the number of samples approached the neighborhood size.

To evaluate whether metric rankings were robust to dataset selection, we assessed their stability across progressively larger subsets of datasets. Rank agreement increased with subset size and plateaued at a median correlation of 0.95 with 10 datasets (Extended Data Fig. 2b), with minimal changes thereafter. This indicates that the 31 datasets and associated annotation frameworks included in our benchmark are sufficient to yield stable and representative metric rankings.

Overall, our benchmarking analysis identified a robust set of ISC metrics that perform consistently well across diverse biological systems, evaluation criteria, and dataset configurations, providing an effective approach for evaluating cell type annotation quality in scRNA-seq data.

### Inter-sample consistency predicts supervised cell type classification performance in the absence of ground truth

Label projection (or label transfer) is a widely used strategy for automated cell type classification in scRNA-seq datasets, in which a pre-annotated dataset serves as a reference to classify cells in new datasets^8,9,16,30,31^. A key limitation of this approach is that evaluating classification performance requires knowledge of the ground-truth cell type labels – which are, by definition, unavailable when applied to new, unannotated datasets. We hypothesized that, if ISC captures cell type annotation quality, it could serve as a fully unsupervised predictor of supervised cell type classification performance.

To test this hypothesis, we compared 14 classifiers across a diverse collection of datasets (Supplementary Table 1). We implemented two complementary evaluation schemes: (i) intra-dataset, using sample-aware train-test splits to assess generalization across biological replicates, and (ii) inter-dataset, training on one dataset and testing on an independent dataset to emulate cross-study label transfer (Fig. 4a). This experiment uses available ground-truth labels in the held-out test, enabling direct assessment of classification performance.

**Figure 4.**
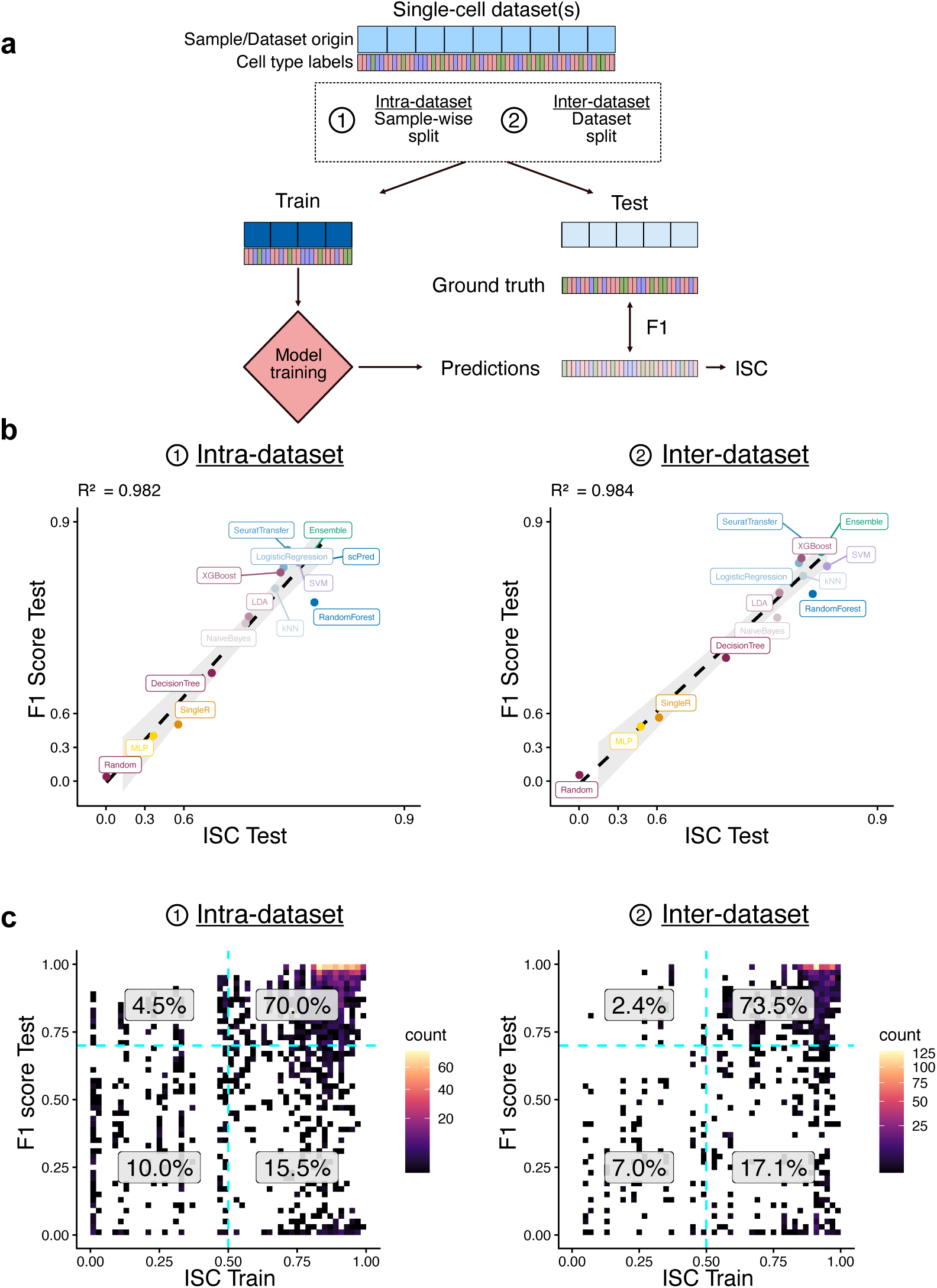
ISC predicts supervised cell type classification performance across multiple datasets and classifiers. (**a**) Schematic of the supervised cell type classification framework. For intra-dataset evaluation, each single-cell RNA-seq dataset was split at the sample level into training and test sets. For inter-dataset evaluation, classifiers were trained on one dataset and tested on a distinct, held-out dataset. (**b**) Relationship between mean F1 score and mean ISC in the test set, shown for intra-dataset (left) and inter-dataset (right) settings. Each point represents the mean per classifier. (**c**) Density plots showing the joint distribution of ISC in the training set (x-axis) and F1 score in the test set (y-axis) for intra-dataset (left) and inter-dataset (right) settings. Density is computed from cell type-level values. Dashed lines indicate thresholds used to define quadrants: ISC = 0.5 and F1 = 0.7. Percentages indicate the proportion of cell types in each quadrant across all datasets. Results are shown for 21 datasets (intra-dataset) and 12 dataset pairs (inter-dataset). Integrated ISC was used, being the product of Task 1-oriented: 2-label silhouette on pseudobulk cosine dissimilarity, and Task 2-oriented: silhouette on RCM dissimilarity. The following classifiers were evaluated: scPred, SVM, RandomForest, SeuratTransfer, LogisticRegression, kNN, NaiveBayes, LDA, XGBoost, DecisionTree, SingleR, MLP, Ensemble, and Random (negative control). Correlations between ISC and F1 were computed per classifier using Pearson correlation; corresponding R² values are reported.

We next quantified supervised classification performance for each cell type using the F1 score and compared these values to per-cell-type ISC, calculated as the product of the two best complementary ISC metrics (task 1 and task 2-oriented, see Methods). Remarkably, average ISC and F1 scores were highly correlated across classifiers in both intra-and inter-dataset settings (R² > 0.98; Fig. 4b), indicating that ISC can accurately predict cell type classification performance in a fully unsupervised manner. This relationship also held at the dataset level (Extended Data Fig. 3a). Correlations were slightly reduced when using individual ISC components alone, supporting the use of the combined ISC metric (Extended Data Fig. 3b).

Interestingly, F1 scores calculated on the test data were also significantly associated with ISC computed on the training data (Fig. 4c), indicating that annotation consistency in the reference is a prerequisite for accurate label transfer. However, high training ISC alone was not sufficient to guarantee strong performance, as some classifiers exhibited low F1 scores even when training annotations were highly consistent (Extended Data Fig. 3c).

Together, these results demonstrate that ISC accurately captures annotation quality and predicts the performance of cell type classifiers without requiring ground-truth labels. These findings establish ISC as a practical criterion for selecting reference atlases and benchmarking automated cell type annotation methods.

### Local and global ISC metrics identify suboptimal cell type annotations and guide their improvement strategy

The previous results revealed complementarity between the top task-1-oriented metric – 2-label silhouette on pseudobulk cosine dissimilarity – and the top task-2-oriented metric – silhouette on RCM dissimilarity. The RCM-based ISC captures cell-type inter-sample replicability: the extent to which a cell type profile can be confidently mapped across samples, making it particularly sensitive to closely related, difficult-to-distinguish cell types. In contrast, the cosine-based ISC captures global cell type compactness, i.e. the cross-sample similarity within a cell type in relation to all other cell types. We refer to these two metrics as local ISC (ISC_L_) and global ISC (ISC_G_), respectively (Fig. 5a).

**Figure 5.**
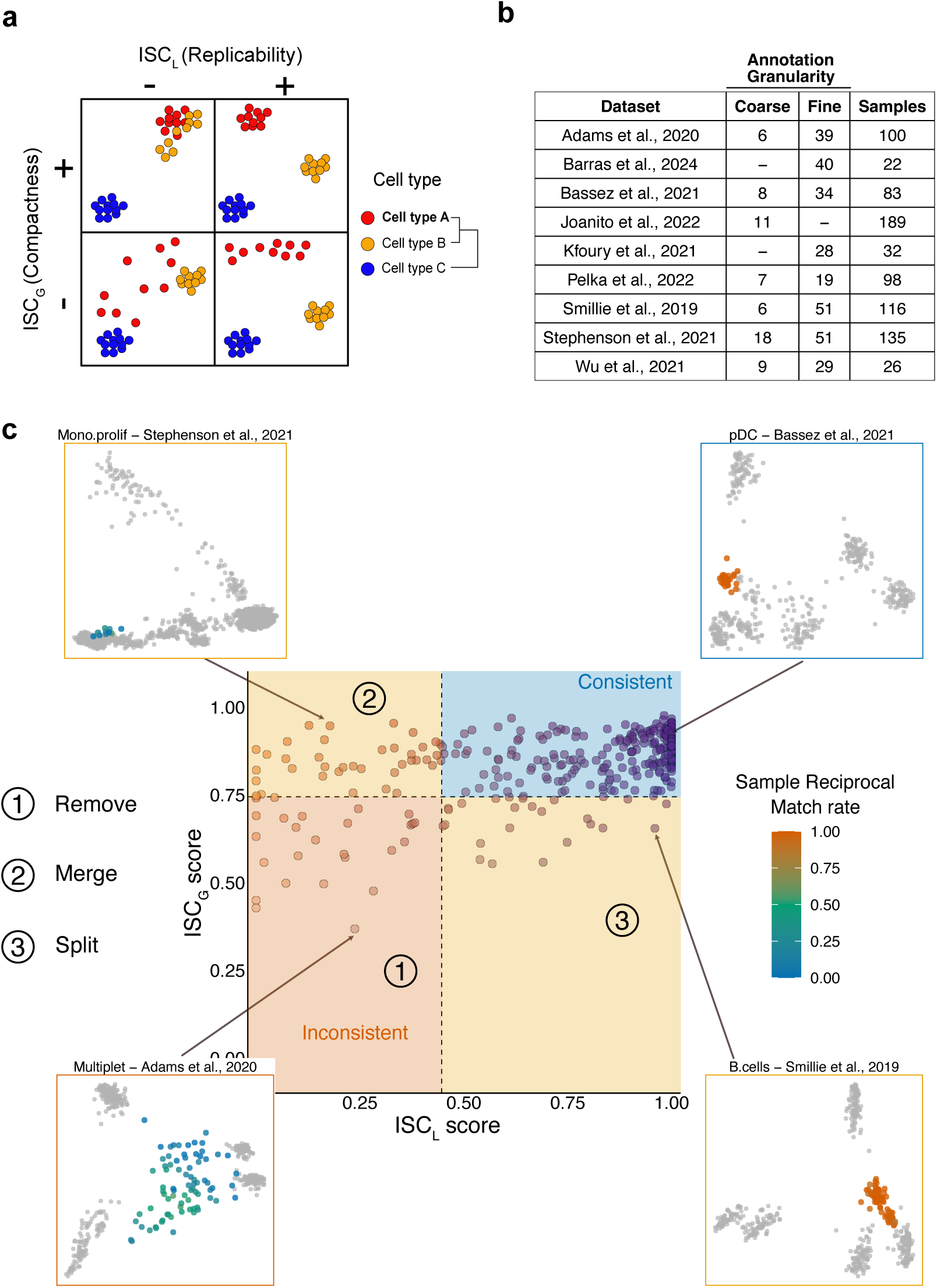
Inter-sample consistency metrics reveal annotation robustness and guide refinement across single-cell datasets. (**a**) Schematic representation of the two-dimensional ISC space. Each dot represents a cell type from a sample in a low dimensional gene expression space. The local consistency (ISC_L_; x-axis) captures cross-sample replicability, whereas the global consistency (ISC_G_; y-axis) captures cross-sample compactness. Each quadrant shows example configurations to illustrate the four combinations of low and high local and global ISC for cell type A. (**b**) Overview of the nine published single-cell datasets included in the meta-analysis, showing the number of biological samples and annotated cell types under both coarse and fine annotation schemes, where available. (**c**) Meta-analysis of author-provided cell type annotations across all datasets. Each point represents a cell type label, positioned by its ISC_L_ and ISC_G_. Dashed lines indicate data-driven thresholds (ISC_L_ = 0.45, ISC_G_ = 0.74; median minus one interquartile range across datasets), partitioning the space into consistent (high–high), inconsistent (low–low), and discordant annotation regimes. Numbered regions suggest annotation actions: (1) low ISC_L_ and low ISC_G_: poor cross-sample consistency, candidate labels for removal or collapse; (2) high ISC_G_ but low ISC_L_: likely over-partitioning, motivating merging of ambiguous or overly fine labels; (3) high ISC_L_ but low ISC_G_: internally heterogeneous populations with reproducible matching but weak global structure, motivating splitting into more coherent subclusters when biologically justified. Insets show representative examples from each regime. For each example, pseudobulk profiles (per sample and cell type) are embedded by PCA as in Fig. 1; the highlighted cell type is colored by its sample-level reciprocal match rate (fraction of other samples in which that cell type is reciprocally matched), while other points from other cell types are shown in grey. Note that this per-sample reciprocal match rate differs from the ISC_L_ metric, which quantifies silhouette scores using reciprocal-classification dissimilarity across all samples for each cell type. ISC_L_: silhouette score on RCM dissimilarity; ISC_G_: 2-label silhouette score on pseudobulk cosine dissimilarity.

To characterize the ranges of ISC_G_ and ISC_L_ observed in real-world atlases and derive empirical guidelines for interpreting ISC scores in practice, we analyzed cell type consistency across nine scRNA-seq datasets spanning diverse tissues and study designs (Fig. 5b and Supplementary Table 1). The distribution of ISC_G_ and ISC_L_ scores revealed distinct consistency regimes (Fig. 5c). Labels with low ISC_G_ and ISC_L_ were enriched for poorly transferable annotations, including multiplets and low-quality labels, which are typically candidates for removal during curation (Extended Data Fig. 4a-b). Labels with high ISC_G_ but low ISC_L_ were highly compact – sharing strong features across samples – but those features lacked specificity and tended to correspond to over-partitioned labels (e.g. proliferating vs. non-proliferating monocytes, Fig. 5c). Conversely, labels with low ISC_G_ but high ISC_L_ lacked strong shared features, yet the available signal was sufficient to correctly match cell type profiles across samples; these may reflect insufficient granularity or highly heterogeneous populations (e.g. a pan B cell population in Fig. 5c). Finally, labels with both high ISC_G_ and high ISC_L_ represent optimal annotations that are at the same time highly replicable and compact (Fig. 5a, 5c, Extended Data Fig. 4a-b).

To facilitate interpretation of ISC_G_ and ISC_L_ across datasets, we derived empirical thresholds from the score distributions obtained in the meta-analysis (median minus one interquartile range for each metric; ISC_G_ = 0.74 and ISC_L_ = 0.45) (Fig. 5c). These thresholds serve as pragmatic guides rather than strict boundaries, partitioning the continuous ISC space into regions associated with distinct annotation refinement strategies: removal of low-quality labels, merging of over-partitioned labels, and subdivision of overly broad populations. Under this framework, most dataset annotations (5 out of 7 at broad granularity, and 7 out of 8 at fine granularity) contained at least one globally or locally inconsistent cell type annotation (Extended Data Fig. 4c). On average, 64% of cell type labels were consistent, whereas 5% showed both low ISC_G_ and ISC_L_, 6% had low ISC_G_, 5% had low ISC_L_, and 3% could not be computed owing to insufficient representation for ISC estimation (fewer than 5 samples and 10 cells per sample).

In summary, this analysis revealed widespread inter-sample inconsistency in cell type definitions across published datasets. Beyond flagging suboptimal annotations, the local-vs-global ISC framework provides a practical guide for improving cell type reproducibility across samples, with each discordant regime pointing to a distinct and actionable refinement strategy.

### ISC-guided refinement of atlas-level cell type annotation

To demonstrate the utility of ISC for identifying and resolving problematic annotations at atlas scale, we applied it to the Human Lung Cell Atlas (HLCA), comprising ∼500,000 cells from 486 individuals across 49 datasets with consensus labels curated by six experts^23^. While most cell type labels in the HLCA showed high ISC_L_ and ISC_G_, we observed a subset with low ISC. These low-ISC labels comprised several respiratory epithelial populations, including club, multiciliated epithelial, and bronchial and nasal goblet cells (Fig. 6a). Inspection of these populations revealed clear clustering by tissue of origin (nose, airway and lung parenchyma) for club and multiciliated cells (Fig. 6b). Expression of canonical marker genes was consistent with such anatomical specialization (e.g. *ELF3* and *FXYD3* were more highly expressed in nose-derived cells, whereas *SCGB1A1* was more highly expressed in airway and lung compartments) (Fig. 6c). These results suggested that transcriptionally and anatomically distinct subpopulations were aggregated under a single label, reducing cell type consistency across samples.

**Figure 6.**
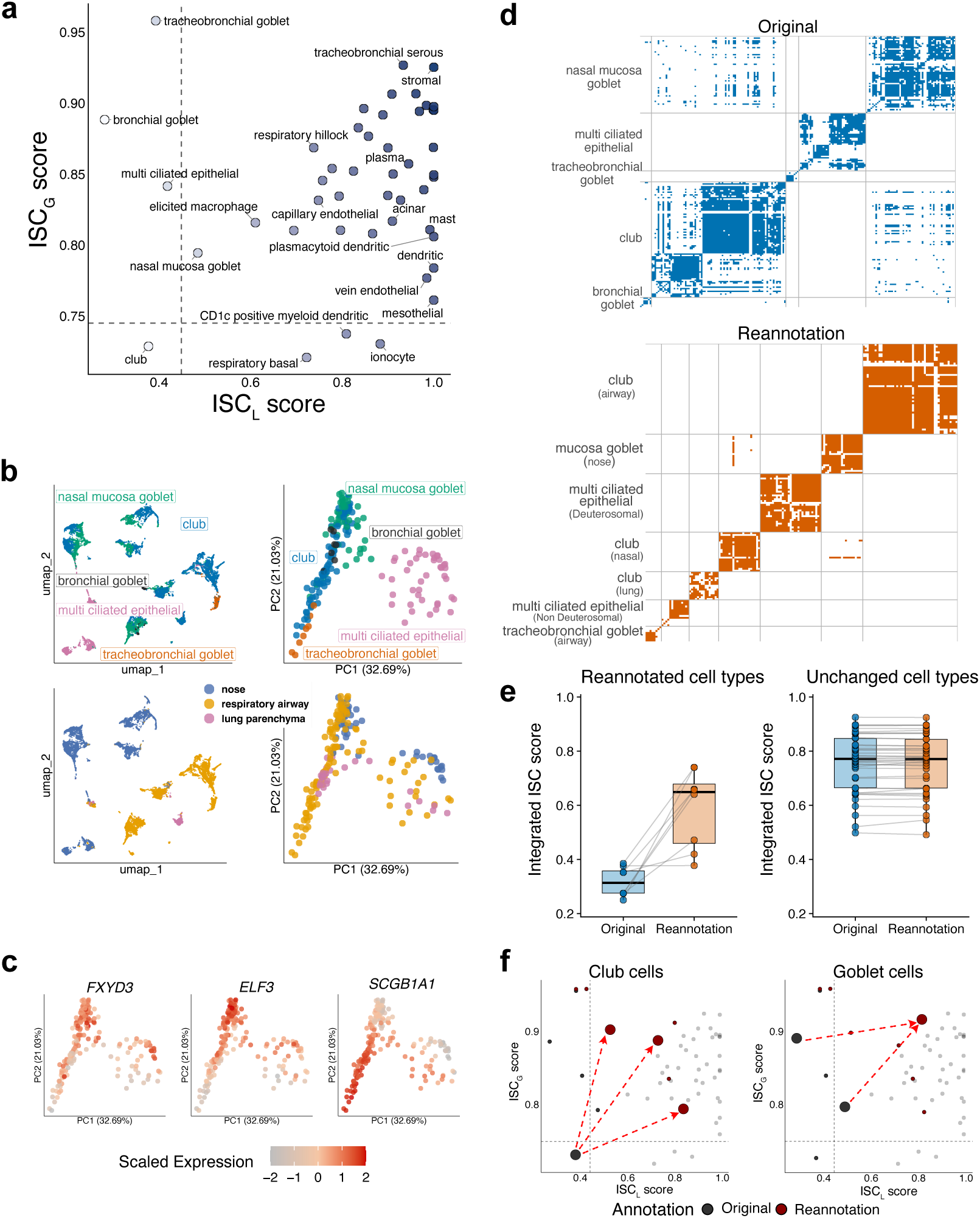
ISC-guided refinement of atlas-level cell type annotation in the Human Lung Cell Atlas. (**a**) Joint distribution of per-cell-type ISC_L_ (silhouette on RCM) and ISC_G_ (2-label silhouette on pseudobulk cosine similarity) across atlas labels (“cell_type”). Each point represents one annotated cell type. Black dashed lines represent empirical thresholds derived from Fig. 5c. **(b)** Low-ISC respiratory epithelial populations visualized in low-dimensional embeddings (bronchial goblet, club, tracheobronchial goblet, multi-ciliated epithelial, and nasal mucosa goblet cells). Left: UMAP of single cells colored by original cell type labels (top) and tissue of origin (bottom). Right: PCA (PC1 vs PC2) of pseudobulk profiles aggregated by cell type and sample, colored by cell type (top) and tissue of origin (bottom). (**c**) Expression of representative marker genes projected onto the same PCA space (PC1–PC2) as in (b, right). (**d**) Reciprocal classification match (RCM) plot for selected low-ISC cell types before (top) and after (bottom) annotation refinement. (**e**) Effect of targeted annotation refinement on integrated ISC (product of ISC_L_ and ISC_G_) for modified labels (left) and unmodified labels (right). (**f**) Embedding of reannotated cell types in ISC_L_-ISC_G_ space as in (a), for Club cells (left) and Goblet cells (right). Each point represents a cell type, selected cell types are enlarged. Grey points denote original annotations while red refined labels. Red dashed arrows indicate the shift in the ISC space following reannotation. Black dashed lines represent empirical thresholds derived from Fig. 5c.

Guided by these observations, we applied targeted refinement of cell type labels, splitting respiratory epithelial cells by tissue of origin. Recomputing ISC under identical preprocessing and metric settings resulted in consistently higher ISC for all modified cell types (Fig. 6d, Fig. 6e left). In particular, club cells, initially characterized by low ISC_L_ and ISC_G_, showed concurrent increases in both metrics upon splitting into tissue-resolved subtypes, reflecting recovery of both inter-sample compactness and replicability (Fig. 6f and Extended Data Fig. 5). In contrast, unmodified labels retained stable ISC values (Fig. 6e right), indicating that refinement selectively corrected problematic annotations without affecting the rest. We also observed that bronchial and nasal goblet cells exhibited high ISC_G_ but low ISC_L_, suggesting over-partitioning. As expected, merging these two labels led to a selective improvement in local consistency (Fig. 6f and Extended Data Fig. 5). In this case, the finer tissue-level cell type partitioning was not supported by the inter-sample consistency criterion. Overall, these results establish ISC as a useful framework to (i) flag heterogeneous or low-quality labels, (ii) diagnose potential underlying causes to guide improvement, and (iii) validate targeted reannotation strategies. This enables the iterative improvement of atlas annotations, increasing consistency while preserving biologically meaningful cell type definitions.

## Discussion

In the context of single-cell transcriptomics, a cell type or cell state is an operational definition rather than a fixed biological entity, reflecting the need to discretize an inherently continuous transcriptional landscape^32–34^. As such, its utility depends on being reproducible, interpretable, and informative across contexts. Thus, a useful cell type label should not only exhibit marker enrichment or within-dataset separability but recapitulate a shared transcriptional identity across independent samples.

In this work, we aimed to formalize this principle through inter-sample consistency (ISC) metrics. By explicitly evaluating cross-sample reproducibility of annotated cell populations, ISC provides a quantitative and interpretable framework for assessing cell type annotation quality in the absence of ground truth labels. Even when the biological function of a cell population is unknown, high ISC provides confidence that it reflects a robust and reproducible biological signal rather than batch- or sample-specific effects.

A few existing methods address a related but different problem: matching cell type labels across multiple studies. For instance, MetaNeighbor^35^ and CellHint^36^ quantify cell type similarity across datasets, identifying correspondences between annotations to support harmonization, integration and meta-analysis. However, they do not explicitly assess cell type annotation quality. Instead, ISC uniquely quantifies consistency across biological replicates within a single study or biological context to evaluate cell type annotation quality. Moreover, ISC provides diagnostic tools to improve cell type annotation, positioned upstream of multi-study harmonization.

Our study identified two major cell type label failure modes that are difficult to diagnose with current validation strategies. Global inconsistency occurs when a labeled cell population is poorly defined in terms of cross-sample compactness and separation from all other cell populations. Local inconsistency, instead, is observed when closely related populations cannot be confidently distinguished across samples. These modes are captured by complementary ISC metrics with distinct representations: global ISC is based on a continuous cosine dissimilarity, whereas local ISC is based on inter-sample binary reciprocal match. Thus, ISC provides a practical framework to guide annotation refinement. Labels with both low global and local consistency are unlikely to be informative and may warrant removal. High global but low local consistency indicates lack of cross-sample replicability and can be fixed by merging conflicting labels. Conversely, high local but low global consistency indicates large inter-sample heterogeneity, potentially fixed by subdivision into finer populations. While domain knowledge remains essential, ISC-guided cell type label refinement can be tested: merging low local consistency labels should increase their local consistency without affecting global consistency, and meaningfully splitting low global consistency labels should increase their global consistency while preserving their local consistency.

ISC metrics have direct implications for the construction of reference atlases and the development of automated cell type classifiers. Low-ISC cell types are unstable targets for supervised classification because they are not replicable across samples. Models trained on such labels are therefore more likely to learn sample-specific boundaries rather than transferable biology. Because ISC predicts classification performance without ground-truth labels, it represents a powerful tool to assess and improve reference quality and to select the best performing models.

A key limitation of ISC is that it requires datasets with multiple biological replicates (here, a minimum of 5) and sufficient cell numbers per label, which may restrict its application in small studies. Although our benchmark indicates that top-performing ISC implementations are robust to typical batch effects, large batch effects may reduce consistency scores. This limitation is less pronounced for RCM-based consistency, which depends on relative cross-sample relationships rather than absolute dissimilarities. A further limitation concerns truly individual-specific cell populations, such as malignant cells, which display highly divergent molecular profiles across individuals, resulting in low consistency scores irrespective of annotation accuracy. Finally, no cutoff defines a universally acceptable ISC score: interpretation remains context dependent and should account for annotation granularity, tissue complexity, and cohort composition.

In summary, ISC provides a framework to evaluate a fundamental yet underappreciated property of cell type annotations: their capacity to generalize across biological replicates. By formalizing inter-sample consistency, distinguishing complementary failure modes, and translating these into actionable refinement steps, ISC helps move single-cell atlas annotation from a largely qualitative practice toward a more quantitative and reproducible process.

## Acknowledgments

This work was partially supported by the Swiss Cancer Research Foundation (grant KFS-5409-08-2021) and the Swiss National Science Foundation (grant 205930).

## Author contributions

All authors conceived and conceptualized the study. J.G. performed software development, analyses, and visualization. J.G. wrote the initial draft. All authors interpreted results and wrote the manuscript. S.J.C. supervised the study. All authors approved the final manuscript.

## Methods

### Datasets

We curated 31 scRNA-seq dataset subsets derived from 17 published studies^23,24,37–39^, spanning eight sample types. These subsets comprised 8 colorectal cancer, 5 blood, 5 healthy lung, 4 basal cell carcinoma, 3 melanoma, 3 healthy airways, 2 breast cancer, and 1 healthy nose datasets. For each subset, Supplementary Table 1 details the study reference, batch, condition, annotation framework(s) evaluated, number of samples retained (range 7–15; median 12), and whether the subset was included in batch-effect or perturbation analyses. These dataset subsets were used for ISC and supervised cell type classification benchmarking.

For the meta-study of ISC metrics, we used some studies mentioned above^24,37^, but also included 6 additional datasets^40–46^. The complete list of datasets, including those used solely for meta-analysis, is provided in Supplementary Table 1.

### Preprocessing

Original study datasets were downsampled to a maximum of 200 cells per sample-label group to balance representation, then split by batch and/or condition where applicable. For each split, we retained at most 15 samples (prioritizing samples with broad label representation and at least 200 cells per sample) and removed splits with fewer than 7 samples.

### Input data type

We computed inter-sample consistency on representations derived from either sample-cell type (i) single-cell distributions or (ii) pseudobulk profiles. Raw counts were log-normalized (Log1p) and filtered to retain cell types present in at least five samples (min_samples = 5), with at least 10 cells per sample and cell type (min_cells = 10). For pseudobulk-based representations, counts were firstly aggregated per sample and cell type and subsequently log-normalized. Unless otherwise specified, computations were carried out on highly variable genes (HVGs; scran-based^47^ selection, 2,000 genes by default, blocking by sample) and, when a low-dimensional representation was used, on a PCA embedding (30 PCs by default).

### Dissimilarity computations

All ISC metrics operate on a cell type-by-cell type dissimilarity matrix in which each annotated cell type from every sample is compared to all annotated cell types across all samples of the dataset, including the same annotated type in different samples. We computed this matrix using three families of dissimilarities:

i. Pseudobulk distances: Euclidean distance, cosine distance (1 − cosine similarity), and Pearson distance (1 − Pearson correlation) between sample-level pseudobulk profiles.
ii. Wasserstein distance: optimal transport between sample-level single-cell distributions (Wasserstein-2; transport-based solver^48^).
iii. Reciprocal classification: bidirectional classification between sample pairs using SingleR^49^. For each pair of samples, one sample was used as the labeled reference and the other as the query, and the procedure was repeated in the opposite direction. SingleR was run in classic mode (de.method = “classic”), in which marker genes are identified by pairwise comparisons among the reference labels and each query profile is scored against every reference label. The best match was defined as the highest-scoring pruned SingleR label. Reciprocal classification was encoded either as a match-based dissimilarity, in which reciprocal matches were assigned low dissimilarity (0) whereas non-reciprocal assignments received higher dissimilarity (1), or as a score-based dissimilarity derived from the paired reciprocal SingleR scores for the two profiles being compared within a 0-1 scale.

### Inter-sample consistency (ISC) metrics

We computed ISC metrics on the dissimilarity matrix described above. Each observation corresponds to one sample-label group. Metrics are computed at the observation level and then aggregated by averaging across all observations sharing the same label, yielding one ISC score per label. Unless otherwise stated, we used the default parameters: k = 5 for neighborhood-based metrics and Ward linkage (Ward D2) for clustering agreement.

#### Silhouette

We computed the silhouette width^50^ of each sample-label observation using the pairwise dissimilarity matrix and the annotation labels as cluster assignments. For an observation $i$, let $a(i)$ be the mean dissimilarity to observations of the same label and $b(i)$ the minimum mean dissimilarity to observations of any other label; the silhouette is

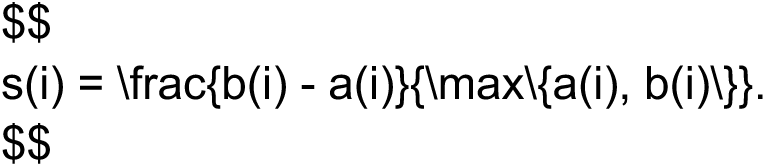

We report the average silhouette width (ASW) per label (range $[-1, 1]$), where higher values indicate tighter within-label cohesion and greater separation from other labels across samples.

#### 2-label silhouette

To specifically quantify how well each label separates from the rest of the annotation, we computed a 2-label silhouette variant. For each observation $i$ with label $\ell$, we define $a(i)$ as the mean dissimilarity to other observations of label $\ell$ and $b(i)$ as the mean dissimilarity to all observations not in $\ell$ (all other labels pooled). The per-observation score is computed as classical silhouette and then averaged per label.

#### Neighborhood purity

We built a K-nearest neighbor (KNN) graph from the dissimilarity matrix and, for each observation, computed the fraction of its $k$ nearest neighbors sharing the same label. By default, $k=5$ and is capped by the maximum number of observations available for any single label. We report the mean neighbor-purity score per label.

#### Clustering agreement

We performed Ward hierarchical clustering (Ward D2) on the dissimilarity matrix and cut the dendrogram into $K$ clusters, where $K$ equals the number of unique labels in the annotation. For each label, we computed the proportion of its observations assigned to its dominant (most frequent) cluster.

#### Nearest-medoid agreement

For each label, we identified a medoid observation that minimizes the within-label sum of dissimilarities (computed from the full symmetric dissimilarity matrix). For each non-medoid observation, we compared its dissimilarity to its own label medoid versus the minimum dissimilarity to any other label’s medoid. The per-label score is the fraction of observations closer to their own medoid than to any other medoid (excluding the medoids themselves).

#### Average similarity

For each observation, we computed (i) the average dissimilarity to other observations with the same label (excluding itself) and (ii) the average dissimilarity to observations of all other labels. We then converted these two averages into a normalized score that increases when within-label dissimilarities are small relative to between-label dissimilarities, implemented as 1 minus the ratio of the within-label average over the sum of within-label and between-label averages.

### ISC Benchmarking

#### Evaluation tasks

We benchmarked ISC metrics on a set of semi-synthetic evaluation tasks designed to probe sensitivity to annotation inconsistencies and robustness to changes in sample composition and experimental conditions:

##### Task 1: Cell type signal degradation

To simulate progressive loss of cell type annotation quality, we introduced controlled label noise by shuffling cell type labels within each sample. For a given perturbation level, a specified fraction of cells was selected at random and reassigned labels drawn from the sample’s original label distribution, thereby preserving per-sample label frequencies. This procedure was repeated across six levels of signal degradation, corresponding to proportions of original labels retained of 1, 0.9, 0.75, 0.5, 0.25, and 0. For each level, three independent replicates were generated.

##### Task 2: Cell type over-partitioning

To simulate unsupported fine-grained annotation, we artificially subdivided a single cell type into two closely related subtypes. For each dataset, the most abundant annotated cell type was selected as the target. Within each sample, cells belonging to this target label were randomly split into two equal groups, with one group reassigned to a new sub-label while the other retained the original label. This procedure was repeated across three independent replicates to account for stochastic variability.

##### Task 3: Granularity

To assess robustness to annotation granularity, we simulated progressively coarser cell type annotations by iteratively merging labels. At each iteration, cell types were represented by expression centroids computed from processed expression values, and pairwise distances between centroids were calculated using Euclidean distance. The two most similar labels were then merged, and ISC metrics were recomputed. This procedure was repeated until only two labels remained, producing an ISC trajectory as a function of the number of retained cell types.

##### Task 4: Cellular complexity

To evaluate robustness to cellular complexity, we systematically varied the number of annotated cell types included in the analysis. For each dataset, we sampled up to 30 subsets of labels spanning a range of sizes, from two labels up to all but one of the original labels. For each subset, cells belonging to excluded labels were removed, and ISC metrics were recomputed on the remaining data.

##### Task 5–6: Number of samples and number of cells per cell type

To assess robustness to cohort size and label abundance, we independently varied the number of samples and the number of cells per cell type. For sample size robustness, we sub-sampled each dataset by retaining decreasing numbers of samples across multiple predefined levels (1, 0.9, 0.7, and 0.5), generating three independent replicates per level. To evaluate robustness to cell type abundance, we downsampled the number of cells belonging to a target label, by default the most abundant label in the dataset, across several levels of retained cell counts proportions (1, 0.75, 0.5, and 0.25). Each downsampling level was evaluated using three independent replicates.

##### Task 7: Batch effects

To evaluate robustness to batch effects, we compared ISC values computed within individual experimental batches to those obtained after combining batches within the same annotation framework.

##### Task 8: Biological perturbations

To assess robustness to biological perturbations, we stratified datasets by biological condition within the same batch and computed ISC separately within each condition. These values were then compared to ISC computed after pooling datasets across conditions.

#### Normalized task performance scoring

For each task in the benchmarking, metric behavior across perturbation levels was summarized using a normalized task performance score from 0 (worst) to 1 (best). ISC values were first averaged across replicates and then summarized per cell type or at the annotation level, depending on the task. For Task 1, where ISC is expected to decrease monotonically as annotation quality declines, performance was quantified with a monotonicity score defined as the fraction of consecutive perturbation steps in which ISC decreased, divided by the total number of steps minus one. For Tasks 3-6, where ISC is expected to remain stable across perturbation levels, robustness was quantified using a constant-fit score defined as 1 minus the normalized root mean squared deviation (nRMSD) of ISC values from their mean. The RMSD was normalized by 0.5, which is the maximum possible RMSD for a metric bounded between 0 and 1, so that the final score spans the full 0 to 1 range. Higher scores indicate flatter, more stable profiles. For Tasks 2, 7, and 8, which involve two discrete conditions, performance was quantified as the consistency drop score: the difference in ISC between the pooled or complete condition and the split condition, clipped to the range 0 to 1. Higher scores indicate that ISC was better preserved under the perturbation.

#### Benchmark ranking and visualization

Benchmarking data were processed as follows: for each evaluated ISC metric, performance scores were computed across eight tasks. Raw results were aggregated and standardized using min-max scaling. Overall score for each ISC metric was calculated as a weighted mean across tasks (weights: 0.2 for signal degradation and over-partitioning; 0.1 for all other tasks). To visualize benchmarking results across multiple tasks and ISC metrics, we used the R package funkyheatmap^51^ (v1.0.0). ISC metrics were ranked by overall score, and task-specific scores were displayed as circles, with size proportional to the median score and color indicating intra-task rank. Resource usage (peak memory and duration) was summarized per method and visualized as bar plots but not used for the overall ranking.

#### Residual effects analysis of consistency metrics and dissimilarity methods

To dissect the contribution of individual consistency metrics and dissimilarity methods across benchmarking tasks, we computed residual effects for each component. For each task, the marginal mean score was calculated for every consistency metric and dissimilarity method by averaging across all methods sharing that component. The residual effect was defined as the difference between this marginal mean and the grand mean score for the task, thus centering effects per task. Residuals were then ranked within each task to facilitate comparison.

#### Stability analysis of ISC metrics rankings across datasets

To assess the robustness of benchmarking method rankings to dataset selection, we implemented a stability analysis. For each group size k (number of datasets), we generated up to 5,000 random pairs of disjoint (non-overlapping) dataset combinations of size k. For each combination, we computed the ranking of ISC metrics based on their weighted mean performance across tasks (using the same task weights as in the main benchmarking analysis). The Spearman correlation between method rankings in each disjoint pair was calculated, quantifying the consistency of rankings across independent dataset subsets.

To summarize how ranking stability changes with group size, we calculated the interquartile range (IQR) of Spearman correlations for each group size. We then modeled the relationship between group size and IQR using a generalized additive model (GAM) with a cubic spline basis (mgcv R package^52^), allowing for smooth interpolation and extrapolation of variability trends. This approach enabled us to estimate the expected reduction in ranking variability as more datasets are included, and to identify the point at which additional datasets yield diminishing returns in stability.

### Supervised cell type classification benchmark

#### Data Splitting

Data splitting was performed in two settings: intra-dataset and inter-dataset. In the intra-dataset setting, samples within each dataset were randomly partitioned at the sample level, with 30% assigned to the reference set and 70% to the query set. This design preserved donor-level structure, batch effects and biological heterogeneity while ensuring complete separation between reference and query samples to prevent data leakage. In the inter-dataset setting, reference and query data were drawn from different datasets matched for annotation reference and biological condition, but differing in batch or study origin. Both transfer directions were evaluated for each eligible dataset pair. In both settings, raw count matrices and per-cell metadata, including sample identity and expert-curated cell type labels, were stored separately for the reference and query data.

#### Data Preprocessing

Unless otherwise specified, classifiers were trained on log-normalized gene expression values obtained by scaling counts to 10,000 per cell followed by log transformation. Highly variable genes (HVG) were identified independently within each dataset based on variance computed from log-normalized expression values. Genes with zero variance were excluded. HVG were defined by ranking genes by variance and retaining the top 2,000 features. All downstream analyses, including feature selection and dimensionality reduction, were restricted to the HVG set. Where applicable, principal component analysis (PCA) was performed on log-normalized HVG expression values after centering (without scaling), retaining up to 30 principal components. Components with zero variance were discarded, and any non-finite values were set to zero.

#### Classification Methods

We evaluated a total of 14 distinct classification algorithms.

##### SingleR (Reference-based Correlation)

SingleR performs per-cell correlation-based classification using the full reference transcriptome. It computes Spearman correlations between query cells and each reference cell, scores each cell type by aggregating correlations, and assigns the cell type with the highest average correlation. Pruned labels (with lower confidence scores filtered) were used for evaluation. No feature selection is performed in SingleR^49^; all genes common to both reference and query are retained.

##### Logistic Regression (Multinomial, L1-regularized)

Multinomial logistic regression was implemented using the R package glmnet^53^ with L1 (LASSO) regularization and alpha = 1. Features were standardized (z-score normalization) before fitting. The optimal regularization parameter (lambda) was selected via 5-fold cross-validation, minimizing classification error. Classes with fewer than five reference cells were excluded to ensure representation across all folds during 5-fold cross-validation.

##### Gradient Boosting (XGBoost)

XGBoost was trained with 100 rounds of boosting using the multi:softprob objective function (probabilistic classification). Hyperparameters were: learning rate (eta) = 0.3, maximum tree depth = 6, subsample = 0.8, and column subsample = 0.8. The model was trained on log-normalized, HVG-selected features without additional dimensionality reduction. The xgboost R package implementing the XGBoost algorithm was used^54^.

##### Multilayer Perceptron (Neural Network)

A feedforward neural network with one hidden layer (25 units) was trained using the nnet package^55^ with backpropagation. Network training was regularized via weight decay (decay = 0.001) and limited to 500 training iterations. To improve convergence and satisfy the independence assumption of backpropagation, features were projected onto principal component space before training.

##### Random Forest

A random forest classifier with 100 decision trees was trained, with each tree allowed to grow to a maximum of 30 terminal nodes, using the R package randomForest^56^. Splitting was performed on HVG-selected log-normalized features without dimensionality reduction.

##### Support Vector Machine (SVM)

SVM was trained using the radial basis function kernel with cost = 1 via the R package e1071^57^. Features were projected onto the first 30 principal components via PCA. Probability estimates were calculated for downstream prediction confidence analysis.

##### K-Nearest Neighbors (kNN)

The kNN classifier assigned each query cell to the most frequent cell type among its k = 15 nearest neighbors in the training set, using the R package class^55^. Euclidean distances were computed in the PCA space.

##### Naive Bayes

Naive Bayes was trained assuming feature independence within each cell type class, using the R package e1071^57^. To better satisfy the independence assumption, features were first reduced via PCA, and the model was trained on these orthogonal principal components. Class conditional distributions were estimated from log-normalized, HVG-selected features. The model assigned query cells to the class with the highest posterior probability.

##### Linear Discriminant Analysis (LDA)

LDA was trained to find linear combinations of features that maximally separate cell type classes using the lda function from the R package MASS^55^. Count data were first log-normalized and HVG-selected, then projected onto the first 30 principal components via PCA with center = TRUE and scale = TRUE (z-score normalization). LDA was then trained on this PCA-reduced feature space, and query predictions were made by projecting query cells into the learned discriminant subspace.

##### Seurat CCA Transfer (Anchor-based Integration)

Seurat’s transfer mapping approach identifies “anchor” cells between reference and query datasets using canonical correlation analysis (CCA), implemented in the R package Seurat^58^. The reference dataset was preprocessed: log-normalized, variable features identified, data scaled and reduced to 30 principal components via PCA using default Seurat parameters. Query data were log-normalized and scaled using reference variable features. Transfer anchors were identified in the first 30 dimensions, and cell type predictions were transferred from reference to query using weighted averaging of anchor-based similarities.

##### Decision Tree

A decision tree classifier was trained using recursive partitioning via the R package rpart^59^ with method “class” and maximum tree depth of 15. Features were projected onto the first 30 principal components via PCA before tree construction.

##### scPred

The R package scPred^60^ implements support vector machine-based prediction with probabilistic outputs. Following scPred guidelines, the reference Seurat object was independently preprocessed: log-normalized, variable features identified, scaled data generated, and PCA performed (30 components). scPred’s getFeatureSpace function extracts a low-dimensional feature space per cell type, and trainModel builds probabilistic models. Query cells were normalized and mapped to this reference feature space using scPredict.

##### Random Classifier (Negative Control)

As a baseline negative control, a random classifier assigned query cells to reference cell types with uniform probability. Random assignments were generated by sampling cell types from the reference with replacement.

##### Ensemble Voting Classifier

An ensemble classifier combined predictions from all 12 dedicated classification methods (excluding Random and Ensemble itself) using majority voting. For each query cell, the cell type receiving the most votes among the 12 independent classifiers was assigned. In case of ties, the first mode in sorted order was selected. Cells on which fewer than one classifier made a valid prediction (all returned NA) were marked as unassigned.

### Data and code availability

All datasets required to reproduce the ISC metrics and supervised cell type classification benchmarking analyses in this study are publicly available through Zenodo at 10.5281/zenodo.18921437.

scTypeEval is available as an R package from BioConductor (https://bioconductor.org/packages/scTypeEval) and GitHub (https://github.com/carmonalab/scTypeEval).

All code used to perform the analyses and generate the results presented in this study is available on GitHub (https://github.com/carmonalab/ISC_benchmark_reproducibility).

