## Extended Data Figures for "Evaluating cell type annotations in single-cell omics in the absence of ground truth"

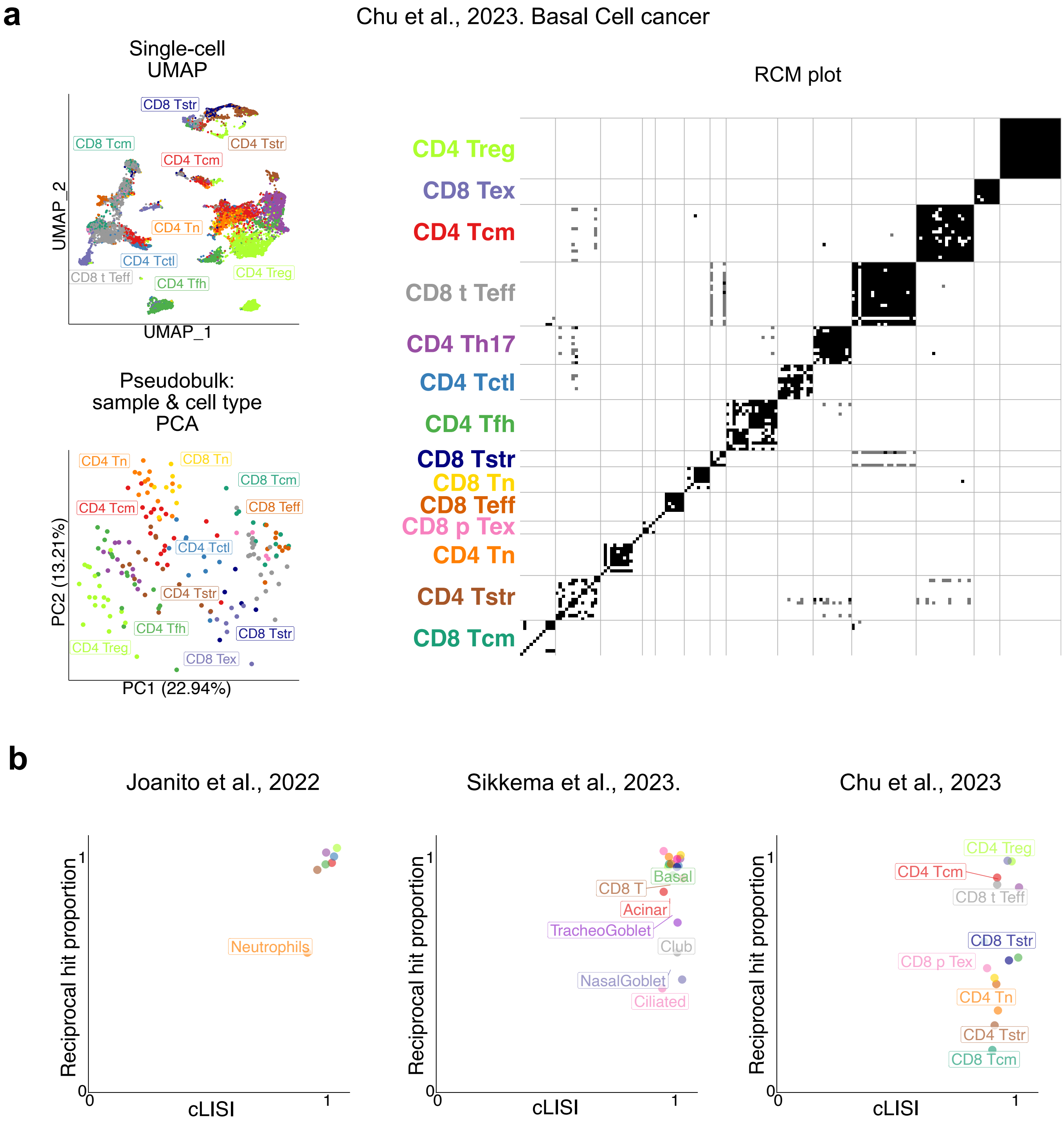


Garnica et al., Extended Data Figure 1

Extended Data Figure 1.

**Inter-sample consistency of expert-curated annotations in carcinoma-infiltrating T cells and comparison with cLISI.** (**a**) Analysis of the Chu et al. 2023 scRNA-seq dataset (22 donors; basal cell carcinoma-infiltrating T cells; original dataset from Yost et al. 2019). The left panels show a UMAP of single-cell transcriptomes colored by annotated cell type and a PCA of pseudobulk gene expression profiles generated by aggregating, within each donor, all cells assigned to the same annotated cell type, yielding one profile per cell type and sample pair. The right panel shows the RCM plot summarizing pairwise reciprocal classification of annotated cell types across all sample pairs, with tile colors indicating reciprocal best match (black), one-directional match only (grey), or no match (white). Symmetrically, each row and column represent a cell type from one sample. (**b**) Comparison between RCM success and cLISI across the Joanito et al. 2022, Sikkema et al. 2023, and Chu et al. 2023 datasets. Scatter plots show, for each annotated cell type, the proportion of sample pairs satisfying RCM for each cell type versus cLISI. cLISI was computed at single-cell resolution with perplexity = 30 and then summarized at the cell type level. Points and labels are colored by cell type, and labels highlight cell types with lower inter-sample consistency, with additional labels shown where space permits.

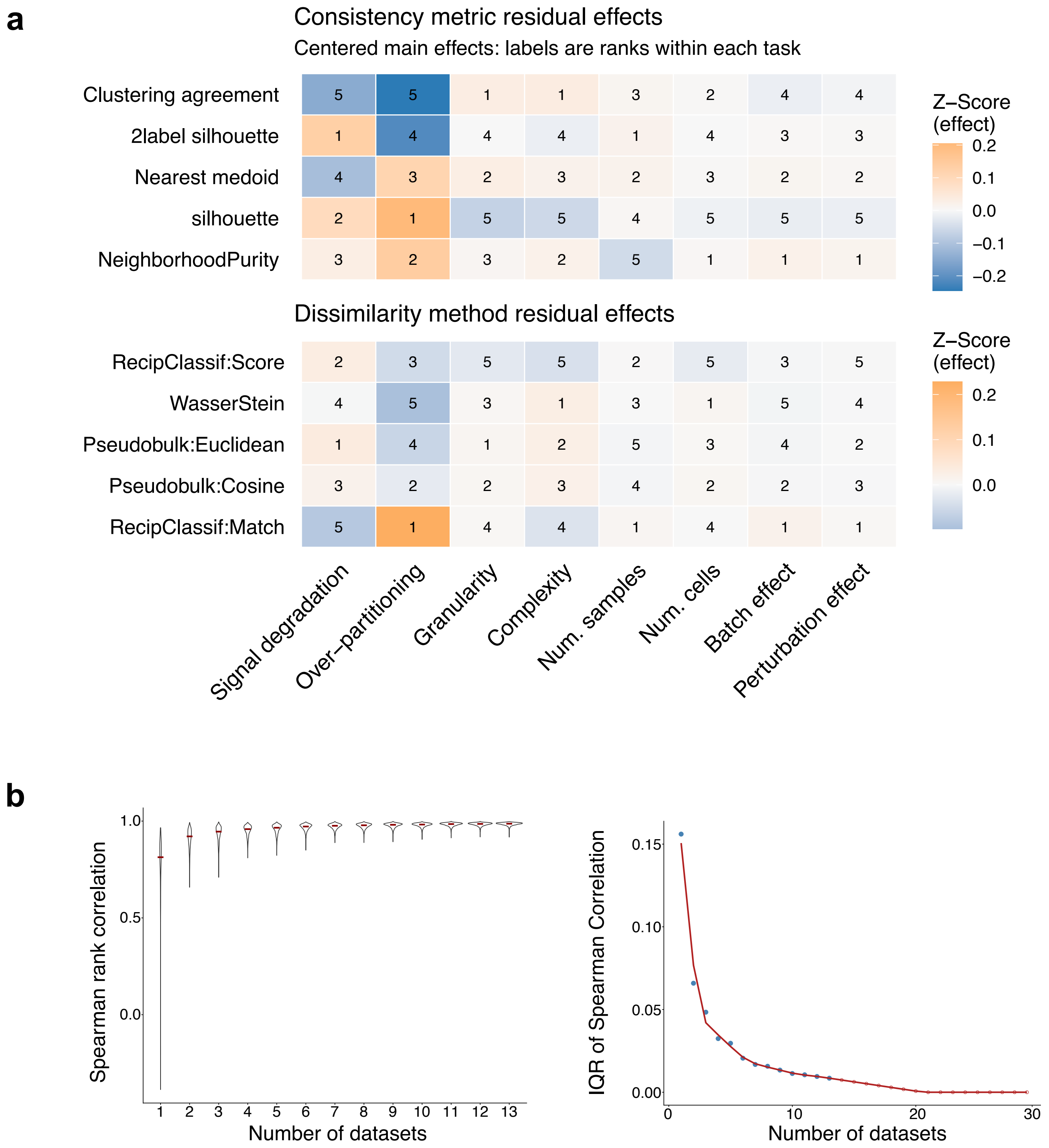


Garnica et al., Extended Data Figure 2

Extended Data Figure 2.

**Component-level effects across benchmark tasks and ranking stability saturation.**
(**a**) Residual-effect decomposition of benchmark performance. Heatmaps show centered main effects (Z-scores) for consistency metrics (top) and dissimilarity methods (bottom) across the eight evaluation tasks. Cell numbers indicate within-task ranks. Warm colors indicate positive residual effects, and cool colors indicate negative residual effects. (**b**) Saturation analysis of ranking stability as a function of dataset number. Left, distribution of Spearman rank correlations between method rankings computed from disjoint dataset subsets at increasing subset size (Number of datasets). Right, interquartile range (IQR) of these correlations versus subset size; blue points indicate observed values, and the red curve shows the fitted trend extended to larger subset sizes by a generalized additive model (GAM). Open red circles indicate extrapolated IQR beyond the observed range.

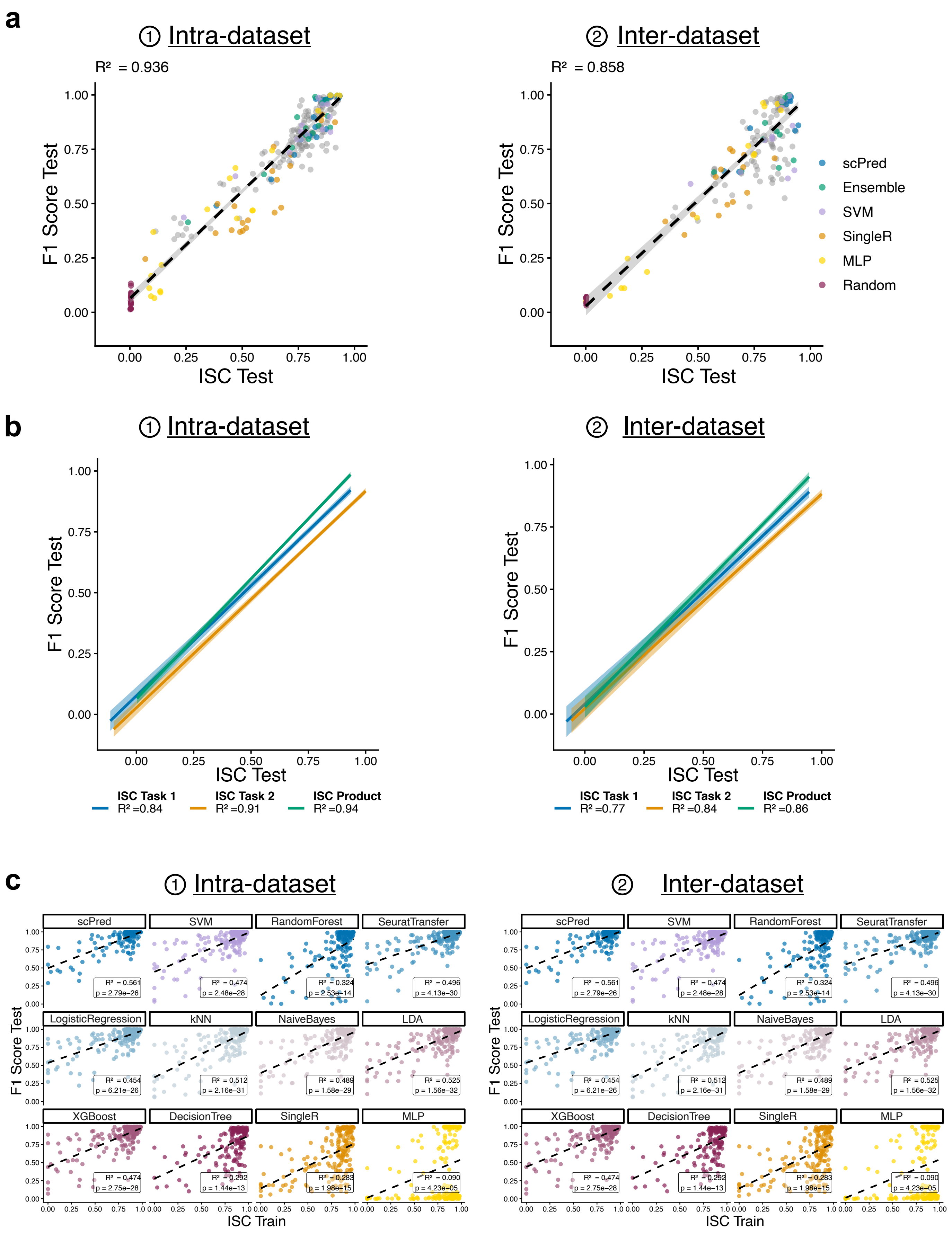


Garnica et al., Extended Data Figure 3

Extended Data Figure 3.

**Robustness of ISC as a predictor of supervised classification performance across classifiers and datasets.**
(**a**) Relationship between mean F1 score and mean ISC (product of local and global) in the test set, shown for intra-dataset (left) and inter-dataset (right) settings. Each point represents the mean per dataset and classifier (as in Fig. 4b, but with dataset-level resolution). (**b**) Correlation between mean F1 score and mean ISC in the test set across intra- (left) and inter-dataset (right) settings. Lines indicate linear regression fits for different ISC variants (ISC Task 1-oriented: 2-label silhouette on pseudobulk cosine dissimilarity; ISC Task 2-oriented: silhouette on RCM dissimilarity, and ISC Product), computed on mean values per dataset-classifier pair. (**c**) Association between ISC (product) computed on the training set (x-axis) and F1 score in the corresponding test set (y-axis), stratified by classifier, excluding Random and Ensemble classifiers, for intra-dataset (left) and inter-dataset (right) settings. Each point represents a cell type. Correlations between ISC and F1 were computed per classifier using Pearson correlation; corresponding $R^2$ and p values are reported in the panels, with multiple testing correction applied using the Benjamini–Hochberg procedure.

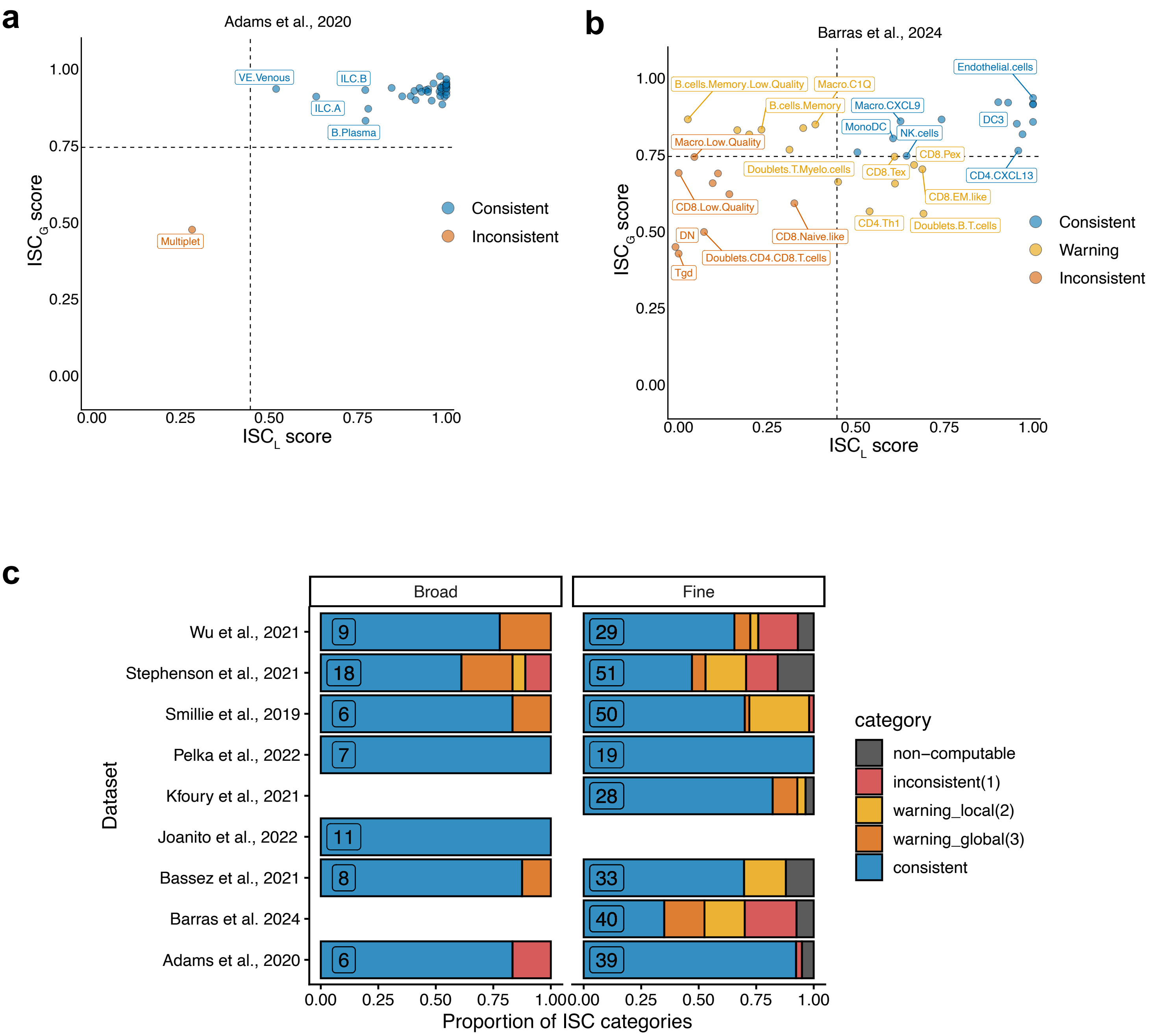


Garnica et al., Extended Data Figure 4

Extended Data Figure 4.

**Dataset-level visualization of local and global inter-sample consistency scores.**

Subset of Fig. 5c for (**a**) Adams et al., 2020 and (**b**) Barras et al., 2024 fine annotations. Each point represents a cell type annotation within the indicated dataset, positioned by its $ISC_L$ score (x-axis) and $ISC_G$ score (y-axis). Dashed lines indicate empirical thresholds: $ISC_L = 0.45$, $ISC_G = 0.74$; median minus one inter-quartile range across datasets, partitioning the space into consistent (blue), warning (yellow), and inconsistent (brown) annotation regimes. Cell types are labeled and colored according to their consistency classification. (**c**) Stacked bar plots showing the proportion of cell types classified into ISC quality categories across datasets in Fig. 5c. Each bar represents a dataset and is stratified by ISC category, including consistent (blue), inconsistent (red), non-computable (dark grey), and warning (yellow) categories. Bars are shown separately for broad and fine annotation levels. The x-axis indicates the proportion of stratified cell types within each dataset. Numbers within each bar indicate the total number of cell types assessed per dataset and annotation level. Non-computable cases reflect cell types for which ISC estimation was not possible due to insufficient cell numbers (minimum 10 cells per sample and 5 samples containing the label).

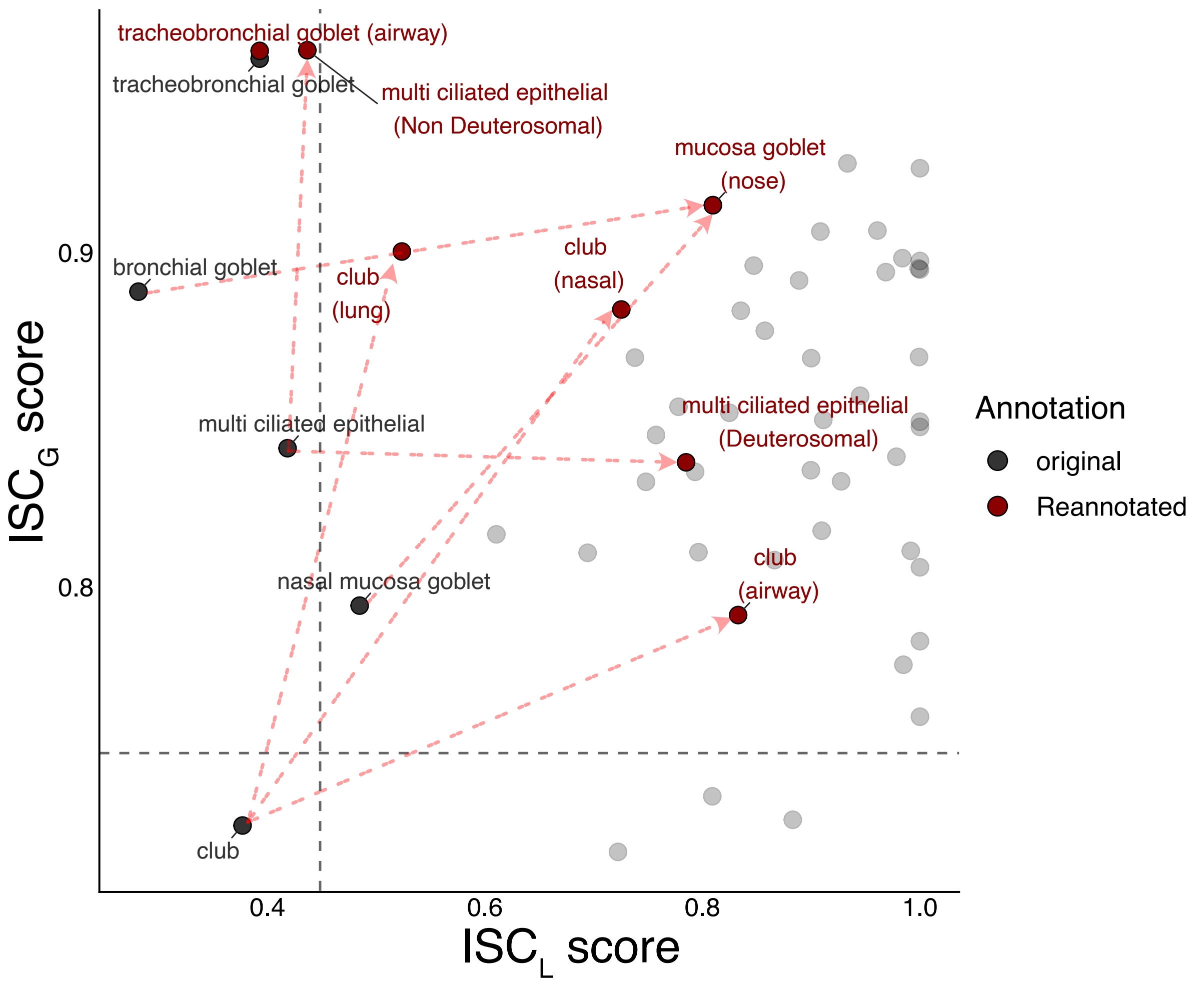


Garnica et al., Extended Data Figure 5

Extended Data Figure 5.

**Global impact of ISC-guided reannotation across refined cell types.**

Embedding of reannotated cell types in $ISC_L$–$ISC_G$ space as in Fig. 6a, shown for all refined labels. Each point represents a cell type, with refined cell types highlighted. Grey points denote original annotations and red points denote refined labels. Red dashed arrows indicate the shift in ISC space following reannotation.
